# Carbamylome analysis reveals regulatory roles for lysine carbamylation in β-cell glycolytic and insulin-processing enzymes

**DOI:** 10.64898/2026.05.29.728846

**Authors:** Youngki You, Hyeyoon Kim, Mereena G. Ushakumary, Marina A. Gritsenko, Hanna E. Walukiewicz, Fangjia Li, Jerry Xu, Ivo Diaz Ludovico, Panshak Dakup, Wei-Jun Qian, Geremy Clair, Gina Many, Raghavendra G. Mirmira, Bobbie-Jo M. Webb-Robertson, Alan J. Wolfe, Christopher V. Rao, Emily K. Sims, Lori Sussel, Ernesto S. Nakayasu

## Abstract

In type 1 diabetes (T1D), insulin-producing β cells are destroyed by an autoimmune response driven by pro-inflammatory cytokines, including interferons. β-cell cytokine signaling is mediated in part by post-translational modifications, such as phosphorylation and acetylation. However, the role of other post-translational modifications in β-cell cytokine signaling represents an important knowledge gap. In the context of autoimmune diseases, lysine carbamylation has gained attention for its role in pathogenesis. Here, we investigate the role of carbamylation in T1D. We found that pancreatic islet cells from the T1D model, non-obese diabetic (NOD) mice, exhibit 11% carbamylation-positive cells, whereas non-diabetic CD1 mice have only 5%. Proteomics analysis of the MIN6 insulin-producing cell line treated with a cocktail of three pro-inflammatory cytokines IFNγ + IL-1β + TNFα identified 284 carbamylated peptides from 222 proteins impacted by the cytokine treatment. Integration of carbamylation and acetylation provided a deep view of the cytokine-regulated PTMs and potential points of interplay. A functional-enrichment analysis revealed that carbamylation was enriched in pathways related to autoimmune diseases, metabolism, DNA replication, and protein translation. Moreover, functional testing demonstrated that carbamylation inhibits the glycolytic enzyme aldolase A and the insulin-processing enzyme carboxypeptidase E, identifying a possible role for cytokine-induced β-cell dysfunction. In summary, protein carbamylation is elevated in islets from NOD mice, and pro-inflammatory cytokine treatment regulates protein carbamylation in MIN6 cells. These data identify carbamylation as a potential regulatory mechanism for β-cell metabolism and insulin production in the context of islet inflammation.

## Introduction

Type 1 diabetes (T1D) results from the autoimmune destruction of insulin-producing β cells, leading to the body’s inability to regulate blood glucose levels. This autoimmune attack is reflected by the production of autoantibodies against proteins of the islets of Langerhans (seroconversion), as well as by the infiltration of immune cells into this tissue (insulitis) [1]. This process is orchestrated at least in part by pro-inflammatory cytokines and chemokines. Even before seroconversion, a circulating transcriptional signature of type I interferon (IFN-α) is detectable in individuals with genetic risk for T1D [2, 3]. This signature is associated with a viral infection, which might contribute to the initiation of the inflammatory response [4]. After seroconversion, type 2 interferon (IFN-γ) is secreted by islet-infiltrating immune cells, leading to the activation of pro-inflammatory signaling [5, 6]. However, this complex cellular signaling is only partially characterized.

Post-translational modifications, such as phosphorylation, deamidation, and ubiquitination, play key roles in autoimmune responses, serving as sources of neoantigens and as signaling transducers [7-9]. Lysine carbamylation (also called carbamoylation or homocitrullination, but distinct from carbonylation or carboxylation) has gained interest in autoimmune disease research [10]. Carbamylation is a non-enzymatic addition of a carbamoyl moiety (–CONH_2_) from cyanate, isocyanate or carbamoyl-phosphate to form N^ε’^-carbamyl-lysine [11]. In the human body, carbamoyl-phosphate is an intermediate of the urea cycle and *de novo* pyrimidine biosynthesis, and its concentration can increase during inflammation [12-15]. Cyanate and isocyanate are formed by urea decay, and it is estimated to reach sub-millimolar concentrations in renal failure [16, 17]. Smoke is another source of cyanate, with concentrations exceeding 100 μM in the body following wildfire exposure [18]. Cyanate is also produced during the oxidation of thiocyanate by the neutrophil enzyme myeloperoxidase [19]. By smoking a single cigarette, a person can inhale 400 µg thiocyanate [20], which can reach 1.6 mM in human saliva [21]. In rheumatoid arthritis, anti-carbamylation autoantibodies have been shown to amplify the pathogenic process [22]. Anti-carbamylation autoantibodies also are produced in other autoimmune diseases, such as systemic lupus erythematosus and Sjögren’s syndrome [22-26]. However, very little is known about the functions of carbamylation in cell signaling.

A previous limitation in studying carbamylation was the lack of unbiased proteomics assays to systematically map and quantify its modification sites. In proteomics, carbamylation poses a challenge because the urea used to denature proteins during sample preparation causes artifactual carbamylation [27, 28]. Carbamylated peptides are also artifacts in acetylomics analysis as they are co-purified with acetylated peptides by anti-acetyllysine antibodies due to their structural similarities [29]. We recently developed a carbamylation analysis pipeline that uses sodium deoxycholate to denature proteins and thus avoid artifactual modification. Then we leveraged carbamylated and acetylated peptide co-purification with anti-acetyllysine antibodies to simultaneously enrich for these peptides [30]. With this method, we studied the roles of carbamylation in bacterial lipopolysaccharide-induced inflammatory signaling in RAW 264.7 macrophage cells. Our analysis identified 2,378 carbamylated proteins, including 360 proteins from diverse functions that were regulated by the lipopolysaccharide treatment. Our study also revealed that carbamylation of linear polyubiquitin chains blocks OTULIN’s anti-inflammatory deubiquitinase activity [30]. Carbamylation also can reduce ubiquitination, but not to the extent of impeding protein targeting to proteasomal degradation [31].

Here, we studied possible roles of protein carbamylation in T1D development. We first investigated carbamylation levels in non-obese diabetic (NOD) mice, a T1D animal model. We also treated the MIN6 insulin-producing cell line with pro-inflammatory cytokines to mimic the insulitis microenvironment during disease development, followed by carbamylome and acetylome analyses. Our results provide a detailed view of the pathways enriched in carbamylated proteins. Moreover, we show the possible regulatory roles of carbamylation in metabolic and insulin-processing enzymes.

## Material and Methods

### Testing the specificity of anti-carbamylation antibodies

The specificity of antibodies was tested by ELISA using carbamylated, acetylated, or citrullinated fetal bovine serum (FBS) prepared in the laboratory. FBS was carbamylated with 1 M potassium cyanate in PBS for 12 h at 37 °C 400 rpm [23]. FBS was acetylated with 5.75% acetic anhydride in 250□mM triethylammonium bicarbonate buffer (pH adjusted to 8.5 with 2 M NaOH) for 1 h at 25 °C with 400 rpm shaking. The reaction was quenched and O⍰acetyl groups cleaved by adding 50% hydroxylamine to 1.85% final concentration and incubating for 30 min at room temperature [32]. FBS was citrullinated with 10 units of peptide-arginine deiminase 4 (SAE0086, Sigma-Aldrich) per mg of protein in 0.1 M Tris-HCl, 5 mM dithiothreitol, and 10 mM CaCl_2_ pH 7.4 for 6 h at 37 °C [33, 34]. Modified FBSs had their buffers exchanged to PBS using Micro Bio-Spin (7326223, Bio-Rad). White polystyrene 96-well plates (NUNC) were sensitized with 10 µg unmodified or modified FBS in 100 µL PBS overnight at room temperature. Plates were washed 3 times with washing buffer (WA126, R&D Systems) and blocked with 1% BSA for 1 h. Primary anti-carbamyllysine antibodies (rabbit polyclonal STA-078 - Cell Biolabs, goat polyclonal conjugated with horse radish peroxidase ab175576 - Abcam, rabbit polyclonal 22428 – Cayman, and mouse monoclonal 23203 – Cayman) were serially diluted in 1% BSA to the concentrations shown in the figures and were loaded and incubated for 2 h at room temperature. Plates were washed 3 times with washing buffer and incubated with secondary anti-rabbit and anti-mouse antibodies conjugated with horseradish peroxidase 2 h at room temperature. Plates were washed 3 times with washing buffer and developed with chemiluminescent reagent (Thermo). Reactions were monitored by luminescence in a plate reader (BioTek, USA).

### Immunofluorescence analysis

Pancreata from 8-9 weeks old female non-obese diabetic (NOD) or CD1 mice were paraffin-embedded and cut into 8-µm thick slices. Paraffin was removed with xylene and tissue was rehydrated with serial dilutions of ethanol before being washed with PBS. Slides were blocked with 4% normal donkey serum (ab7475, Abcam) in PBS containing 0.1% Triton X-100 (PBS-T) for 2 h. After blocking, slides were incubated overnight at 4 °C with 1:500 diluted anti-insulin antibody (3014; Cell Signaling Technology) and 1:200 diluted monoclonal anti-carbamylation antibody, then washed with PBS-T. Fluorescent secondary antibodies (1:200 dilution) and DAPI were incubated in the blocking solution for 1 h at room temperature. After washing with PBS-T, slides were mounted with Prolong Gold (P10144, Thermo) and imaged on an epifluorescence microscope (Leica).

### MIN6 β cell line culture and treatment

MIN6 mouse insulinoma cell line was purchased from Millipore Sigma and cultured in DMEM supplemented with 10% FBS, 50 μM 2-mercaptoethanol, and 1% penicillin-streptomycin at 37 ºC under 5% CO_2_ atmosphere. Cells were treated for three time points to study early and late signaling events, 15 min, 2 h, and 24 h with 100 ng/mL IFN-γ (485-MI-100, R&D Systems) + 10 ng/mL TNF-α (410-MT-010, R&D Systems) + 5 ng/mL IL-1β (401-ML-005, R&D Systems), a combination that induces optimal inflammatory signaling based on our previous work [35].

### Protein digestion, labeling and peptide enrichment

Cell pellets were lysed in 50 mM triethylammonium bicarbonate buffer containing 12 mM sodium deoxycholate at 4 ºC for 15 min followed by 95 ºC for 5 min. Protein concentration was measured by BCA assay (Thermo Fisher Scientific). Cell lysates were reduced with 10 mM dithiothreitol at room temperature for 30 min, and cysteine residues were alkylated with 50 mM iodoacetamide in the dark at room temperature for 30 min. The reaction was diluted 5-fold with 50 mM triethylammonium bicarbonate buffer and digested with 1:25 trypsin:protein ratio and 1:50 endoproteinase Lys-C/protein ratio overnight at room temperature. Enzyme reactions were stopped by adding trifluoroacetic acid (0.5% final concentration). Peptides were desalted by solid phase extraction using C18 cartridges (Phenomenex) and dried in a vacuum centrifuge. To normalize the loading for tandem mass tag (TMT) labeling, peptides were resuspended in water, quantified by BCA assay, and equal amounts were aliquoted and dried in a vacuum centrifuge. Labeling was done with an optimized ratio of TMT to peptide amount of 1:1 (w/w) [36], and samples were fractionated by high pH reverse phase chromatography on a reversed phase XBridge BEH C18 (250 mm × 4.6 mm column containing 3.5-μm particles) (Waters, 186003943) using Agilent 1200 HPLC System and concatenated into 6 fractions. Fractions were first subjected to phosphopeptide enrichment using an immobilized metal affinity chromatography (IMAC) method. IMAC phosphopeptide enrichment was performed using AssayMap Fe (III)-NTA cartridges on the Agilent AssayMap BRAVO Sample Prep platform with the Agilent AssayMap Phosphopeptide Enrichment v.2.1 protocol [37]. Phosphopeptides were saved for future analyses. Samples were further concatenated into 4 fractions, and carbamylated and acetylated peptides were enriched from the IMAC flow-through with anti-acetyllysine antibodies using PTMScan® Acetyl-Lysine MotifMagnetic Immunoaffinity Beads (99064, Cell Signaling Technology) and protocol developed for KingFisher magnetic particle processor [38].

### Mass spectrometry and data analysis

Peptides enriched with anti-acetyllysine antibodies were dissolved in 2% acetonitrile and 0.1% trifluoroacetic acid were separated using a reversed-phase column (packed in-house into a 25-cm length of 360 µm o.d. x 75 µm i.d. New Objective fused silica picofrit tubing with ReproSil-Pur 120 C18-AQ 1.9 µm beads) connected to a nanoACQUITY UPLC system (Waters). The column was set to 50 °C using an AgileSLEEVE column heater (Analytical Sales and Services). Peptides were eluted with a linear gradient from 8% to 35% acetonitrile in 0.1% formic acid over 100 min at a flow rate of 200 nL/min. MS data were collected with an Orbitrap Fusion Lumos mass spectrometer (ThermoFisher Scientific). Precursor spectra were collected from 350-1800 m/z at a resolution of 60K along with data-dependent HCD MS/MS spectra at a resolution of 50K. Precursor ions with charges 2+ to 6+ were isolated for MS/MS at a width of 0.7 m/z and fragmented using a normalized collision energy of 30%, and dynamically excluded for 45 s. Data were processed with MSFragger (v 4.1) [39] using FragPipe (v21.1) by matching against the mouse reference proteome database from Uniprot Knowledgebase (downloaded on July 7, 2024). Searching parameters included protein N-terminal acetylation, oxidation of methionine, lysine acetylation and carbamylation as variable modifications, and carbamidomethylation of cysteine residues and tandem mass tags on peptide N-termini and lysine residues as fixed modification. Enzymatic cleavage was set to specific trypsin digestion in both termini. Peptide and fragment mass tolerances were set at 4.5 and 20 ppm, respectively. For re-analyzing the human single islet nanoPOTS proteomics data [40], the same parameters were used except that the human reference proteome from Uniprot Knowledgebase was used as the database, and that no TMT labeling was used in these samples.

### Statistical and pathway analysis

Reporter ion intensity values were normalized by median centering before submitting to Student’s *t*-test using Perseus v.2.0.3.1 [41]. Functional-enrichment analysis was performed using the Database for Annotation, Visualization and Integrated Discovery (DAVID) [42] with the KEGG annotation.

### Aldolase A assay

Recombinant human aldolase A (BLC-01250P, Beta Lifescience) was carbamylated and had its buffer exchanged to PBS as described above. As a control, we performed a mock incubation of an aldolase A aliquot in PBS alone without potassium cyanate. Protein carbamylation was confirmed using Western blot using mouse a monoclonal anti-carbamylation antibody (23203, Cayman) as previously described [43]. Aldolase activity was measured with a colorimetric assay kit (ab196994, Abcam) with absorbance at 450 nm using a plate reader (Tecan M100).

### Carboxypeptidase E (CPE) assay

Recombinant human CPE (10069-H08H, Sino Biological) was carbamylated, had its buffer exchanged with PBS, and confirmed by Western blot as described above. CPE activity was measured by fluorescence using BenzoylAlaArgOH (G4145, Bachem) as substrate and *o*-phthaldialdehyde (P0657, Sigma), which becomes fluorescent upon binding to the arginine released by the enzyme. A total of 75 ng was incubated with 0.5 mM BenzoylAlaArgOH in 50 mM sodium acetate, pH 5.5, containing 5 μM ZnCl_2_ for 10 min at room temperature with shaking at 600 rpm. The reaction was terminated by adding 150 µL of 15 mM *o*-phthaldialdehyde in 0.2 M NaOH containing 0.1% mercaptoethanol (v/v). A total of 200 μL of each sample was transferred into black 96-well plates and measured by fluorescence on a plate reader (M2e, Molecular Devices) with excitation and emission wavelengths of 330 nm and 450 nm, respectively. Specific activity was calculated with the following formula: (adjusted fluorescence [discounting background] x conversion factor [pmol/fluorescence unit) / (incubation time [min] x amount of enzyme [μg]).

## Results

### Elevated protein carbamylation in pancreas of non-obese diabetic (NOD) mice

We investigated protein carbamylation levels in the pancreas of diabetic NOD mice vs. non-diabetic CD1 mice, both at 8-9 weeks old, using immunofluorescence microscopy. At this age, NOD mice have autoimmune responses and insulitis but have not yet developed hyperglycemia. As anti-carbamylation antibodies might cross-react with acetylation and citrullination due to their structural similarities to carbamylation, we first tested 4 commercial antibodies for their specificity using ELISA. A mouse monoclonal antibody showed the most specificity (**Supplemental text and Fig. S1**) and was used to stain the mouse pancreas slices alongside an anti-insulin antibody to stain β cells and DAPI to stain nuclei. A noticeable higher level of protein carbamylation was observed in pancreata from NOD mice compared to CD1 mice (**Fig. S2**). Insulitis in NOD mice also was characterized by lymphocyte infiltration, as evidenced by the high nuclear density (depicted by DAPI stain) near the islets (**Fig. S2**). To have a quantitative measurement, we counted the number of carbamylation-positive cells in islets. In CD1 mice, 5.3% of the islet cells were carbamylation-positive, whereas in NOD mice, 11.0% of the cells were carbamylation-positive (**Fig. 1**). In summary, these results indicate a higher level of protein carbamylation in the pancreas and islets of NOD compared to CD1 mice.

**Figure 1.**
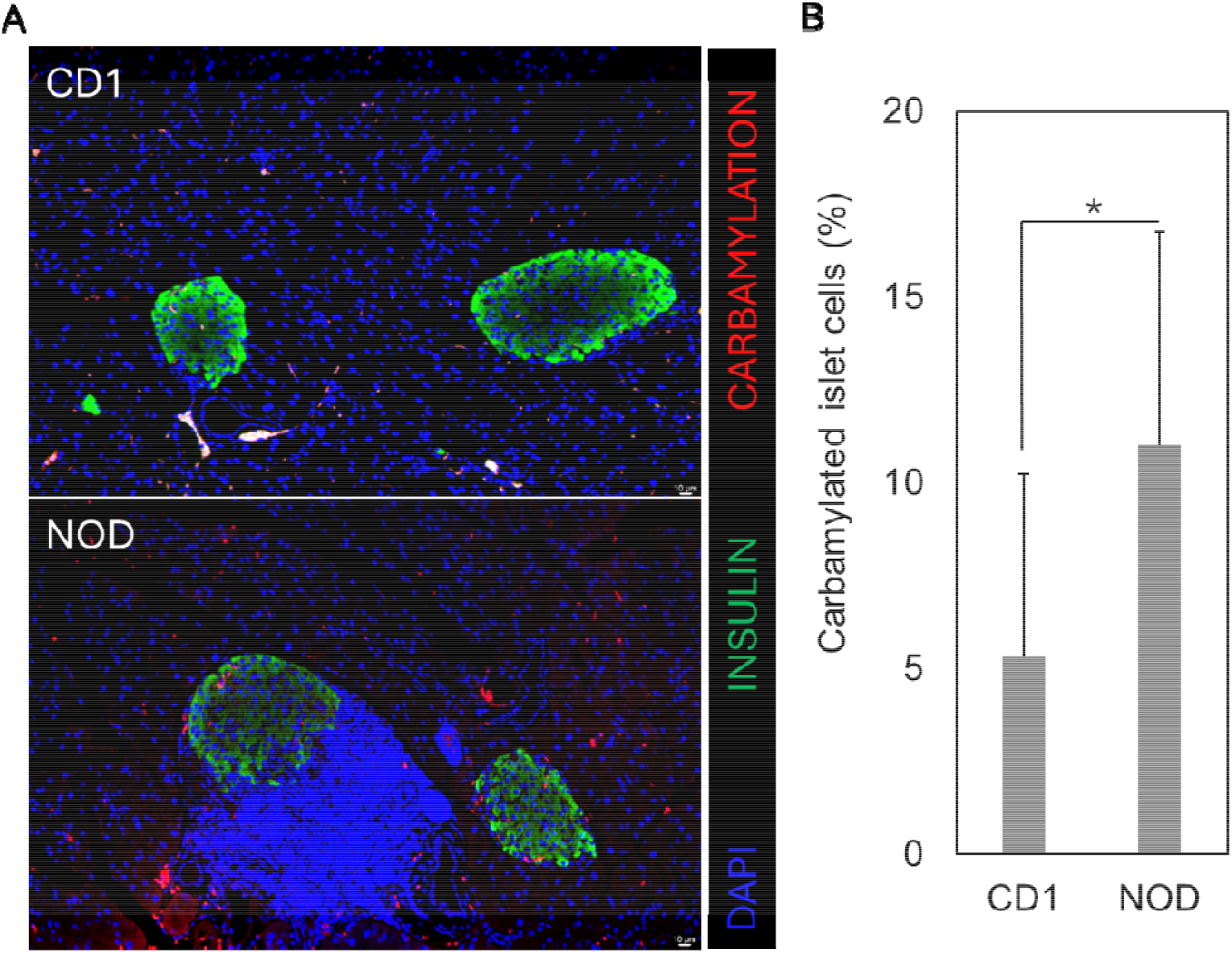
Lysine carbamylation profile of pancreata from non-diabetic CD1 and non-obese diabetic (NOD) mice. (A) Pancreata (n=5) from 8-9-weeks old mice were immunostained for carbamylation, insulin and nucleic acid (DAPI) and imaged in a fluorescence microscope. In NOD mice tissue, there is a noticeable infiltration of lymphocytes, marked by the high density of nuclei (DAPI staining). (B) Quantification of cells carbamylation stain positive cells within islets. * p≤0.05 by Student’s *t*-test.

### Carbamylated proteins in Min6 cells

To study possible functions of lysine carbamylation during insulitis, we treated the MIN6 murine insulin-producing cell line with a cocktail of three pro-inflammatory cytokines, composed of IFNγ + IL-1β + TNFα, for 15 min, 2 h, and 24 h and submitted to carbamylation/acetylation analysis (**Fig. 2A**). A total of 607 carbamylated peptides from 392 proteins were identified. In addition, we identified 12,141 acetylated peptides from 2,841 proteins. From these peptides, 284 carbamylated peptides and 5,860 acetylated peptides were regulated in response to the cytokine treatment (**Figure 2A-B, Tables S1-S2**). Several of the carbamylated proteins with the highest regulation in response to the cytokine treatment were similarly regulated by acetylation, including the histones H2B and H4, RNA exonuclease Rexo1, heat shock protein Hspa9 and the histone acetyltransferase p300 (**Figure 2A-B**). We performed a functional enrichment analysis to identify pathways potentially regulated by carbamylation. A total of 48 pathways were enriched among the differentially carbamylated proteins (**Table S3**). Combining carbamylation and acetylation increased the number of enriched pathways to 104 (**Table S4**), being 41 in common to the carbamylation-enriched pathways alone (**Figure 2C**). Among the 41 common pathways, the average number of proteins per pathway increased 4.7-fold (**Figure 2D**), providing a deeper view of the pathways regulated by the cytokine treatment. Overall, differentially carbamylated proteins were enriched in pathways related to autoimmune diseases, cellular metabolism, DNA replication and gene expression, and protein synthesis and processing proteins (**Figure 2E**), as discussed in the following sections.

**Figure 2.**
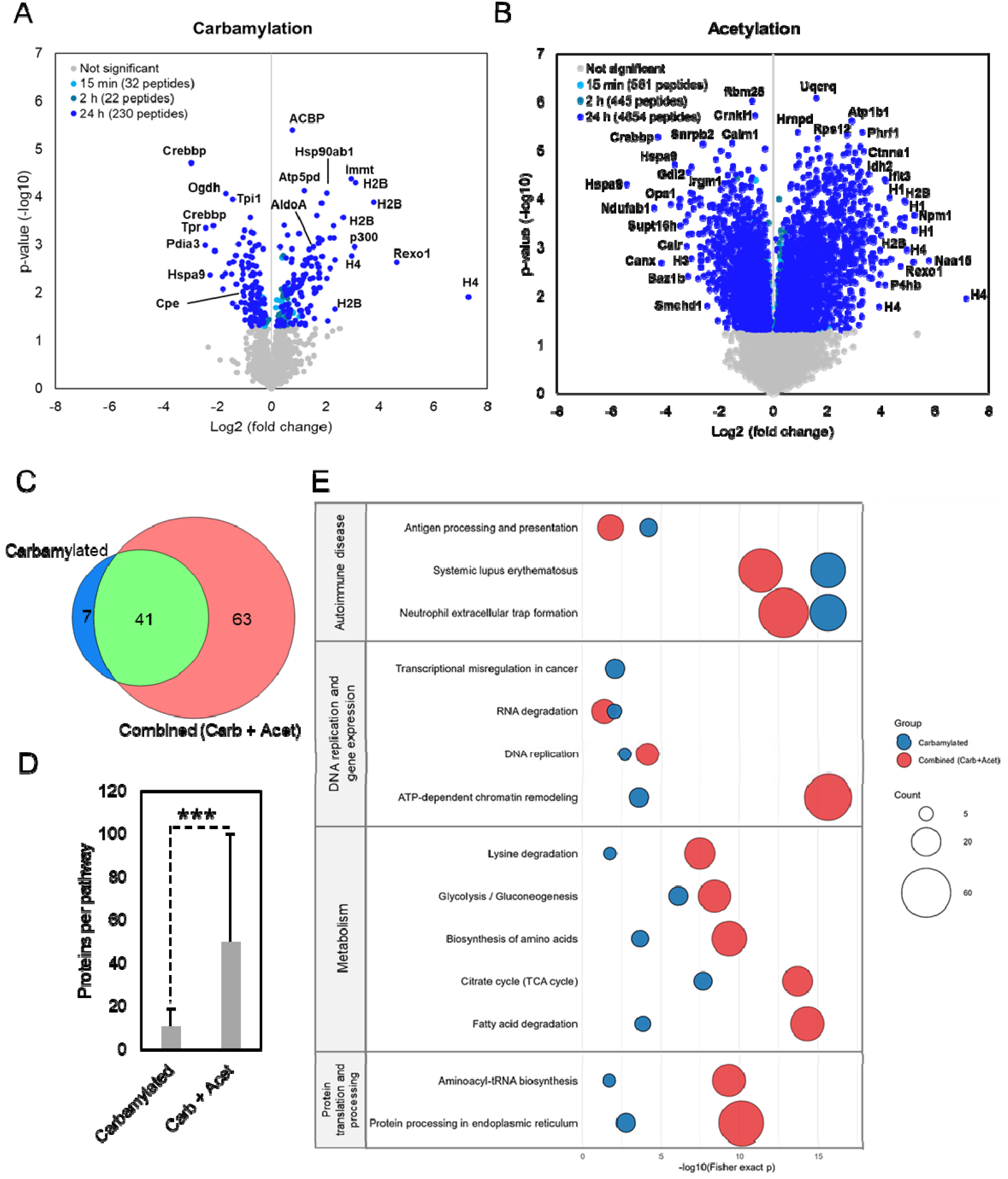
Carbamylome/acetylome analysis of MIN6 insulin-producing cells with pro-inflammatory cytokines. MIN6 cells (n=3) were treated with a cocktail of cytokines (IFNγ + IL-1β + TNFα) for 15 min, 2 h and 24 h, and submitted to carbamylome/acetylome analysis. Volcano plots of quantified carbamylated (**A**) and acetylated peptides (**B**). (**C**) Venn diagram of the number of pathways enriched in differentially regulated carbamylated vs. combined carbamylated and acetylated proteins. (**D**) Number of proteins per pathway. *** p≤0.001 by Student’s *t*-test. (**E**)Selected pathways enriched in differentially carbamylated (blue circles) and carbamylated + acetylated proteins (red circles). The number of carbamylated or acetylated proteins is represented by the circle sizes. The complete lists of enriched pathways are listed in Tables S3-S4.

### Carbamylation enrichment on autoimmune diseases related pathways

Three of the enriched pathways, including the top 2 most enriched pathways, Neutrophil extracellular trap formation and Systemic lupus erythematosus, were related to autoimmune diseases. The third pathway, related to autoimmune diseases, was Antigen Processing and Presentation (**Figure 2E**). In this pathway, 6 proteins were differentially carbamylated with the cytokine treatment, including the heat shock protein Hsp90 and the protein disulfide isomerase Pdia3, which are carbamylated at multiple time points and multiple sites (**Figure 3**). The remaining proteins were calnexin (Canx), calreticulin (Calr), heat shock protein Bip, and transcription factor Rgx5 with a single regulated carbamylated site each (**Figure 3**). The inclusion of acetylation data provided a deeper view of the proteins in this pathway that are modified in response to cytokine treatment, with 13 differentially acetylated proteins (**Figure 3**). In summary, the Antigen Processing and Presentation pathway is highly modified via carbamylation and acetylation in response to the cytokine treatment.

**Figure 3.**
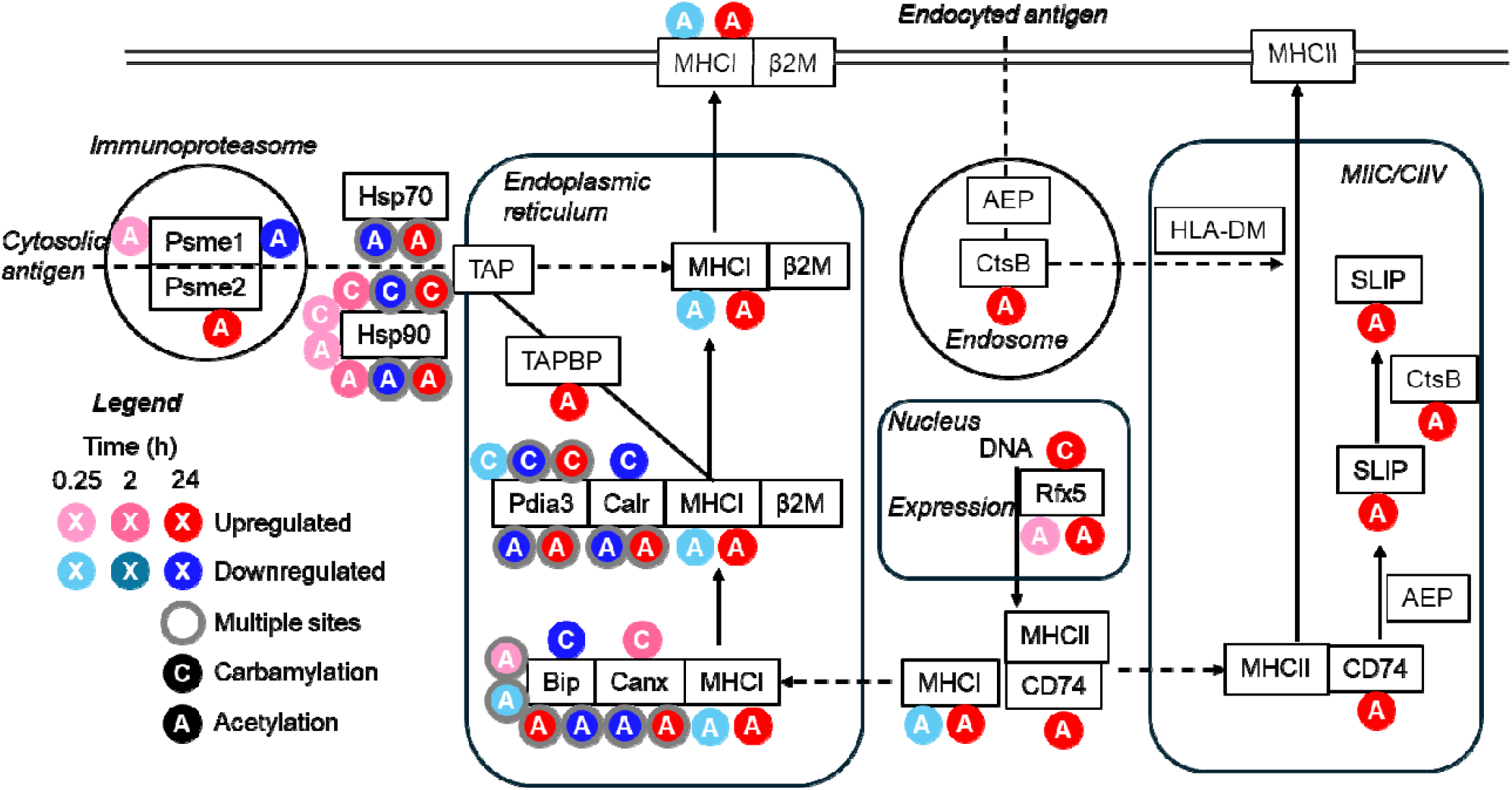
A map of the differentially carbamylated and acetylated proteins of the Antigen processing and presentation pathway.

### Carbamylation enrichment on pathways related to DNA replication and gene expression

Differentially carbamylated proteins also were enriched in pathways related to DNA replication and gene expression, including ATP-dependent chromatin remodeling, DNA replication, transcription misregulation in cancer, and RNA degradation (**Figure 2E**). The pathway with the highest coverage among them is the ATP-dependent chromatin remodeling with 9 and 55 proteins with differential carbamylation and acetylation, respectively (**Tables S3-S4**). A simplified portion of this pathway is represented in **Figure 4**. There is a noticeable overlap of carbamylation and acetylation sites on histones. For instance, 6 out of the 14 histone carbamylation sites had similar temporal regulations as acetylation, 4 had opposite regulations and two had shifts in the temporal regulation (**Figure 4**). We also observed carbamylation of histone acetyltransferases, representing a possible point of crosstalk between lysine carbamylation and acetylation. CREBBP protein, which acetylates histone 3 at lysine 27, has its carbamylation levels at lysines 1584 and 1745 reduced 24 h post-cytokine treatment (**Figure 4**). Histone acetyltransferase p300, which acetylates histones H3 and H4 at multiple sites, has its carbamylation at lysine 1541 increased 24 h post-cytokine treatment (**Figure 4**). These results show that carbamylation differentially modifies chromatin proteins in response to cytokine treatment and may crosstalk with acetylation.

**Figure 4.**
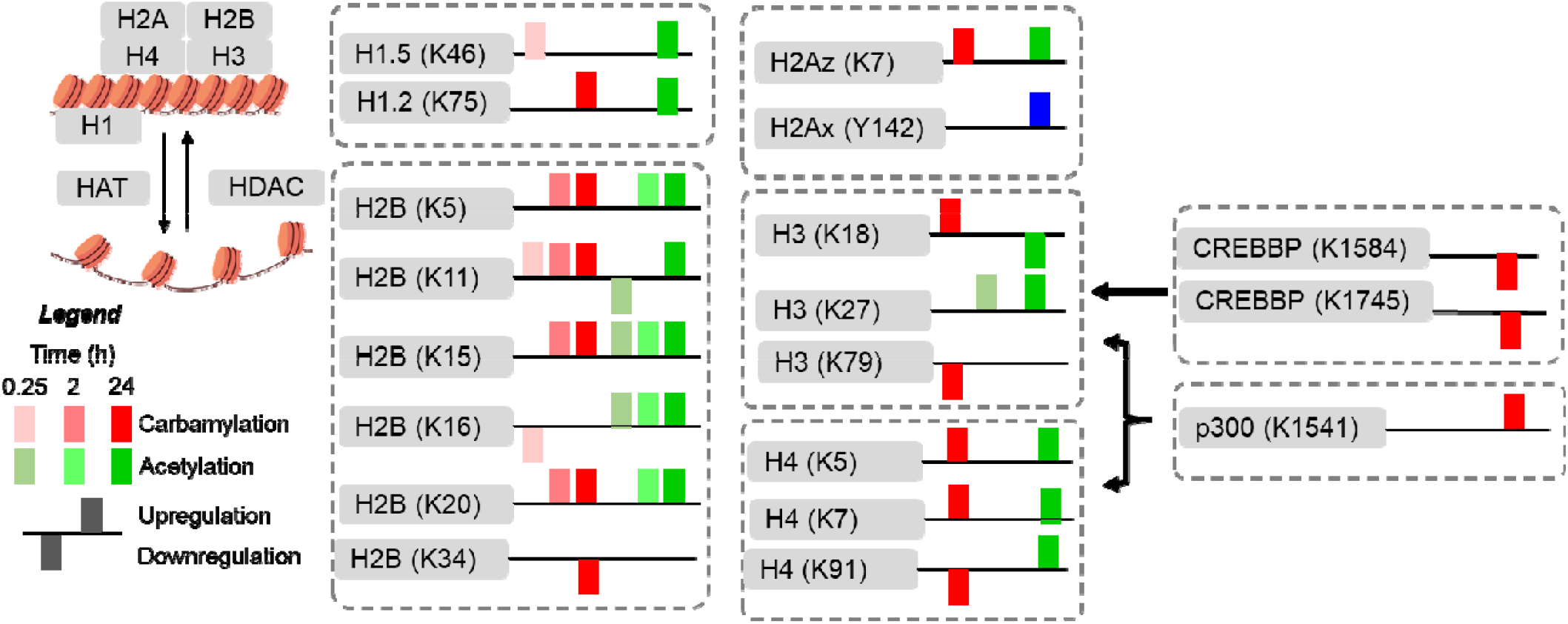
A simplified map of the ATP-dependent chromatin remodeling pathway.

### Differential carbamylation and acetylation of metabolic proteins

A total of 17 out of the 48 pathways enriched in differentially carbamylated proteins were related to metabolic processes (**Figure 2E, Table S3**). Central carbon metabolism was one of the most affected processes. Combining Glycolysis/Gluconeogenesis with Fatty Acid Degradation and TCA cycle, 15 enzymes were differentially carbamylated in response to the cytokine treatment (**Figure 5A**). When including acetylation, almost all the enzymes of these pathways were differentially modified in response to the cytokine treatment (**Figure 5A**). One example is aldolase A (Aldoa), which is differentially acetylated at all 3 time points (**Figure 5A**). In addition, carbamylation of aldolase A at lysine 108 was 4-fold increased 24-h post cytokine treatment (**Figure 5B**). The crystal structure of the human aldolase A (PDB ID 6XMH) shows that this lysine residue is situated close to the active site, having an electrostatic bond to tyrosine 364, a residue that promotes substrate specificity by accommodating the substrate dihydroxyacetone phosphate (**Figure 5C-D**) [44]. Carbamylation of lysine 108 would not only affect the electrostatic interaction with tyrosine 364 but also occludes the substrate binding site by interfering with lysine 147 (**Figure 5E**). To test if lysine carbamylation affects aldolase A activity, we pan-carbamylated a human recombinant aldolase A with potassium cyanate and measured its activity. Carbamylation reduced aldolase A activity to trace levels (**Figure 5F-G**). These results show that carbamylation sites regulated by the cytokine treatment are enriched among the proteins of the central carbon metabolism and can regulate their enzymatic activity.

**Figure 5.**
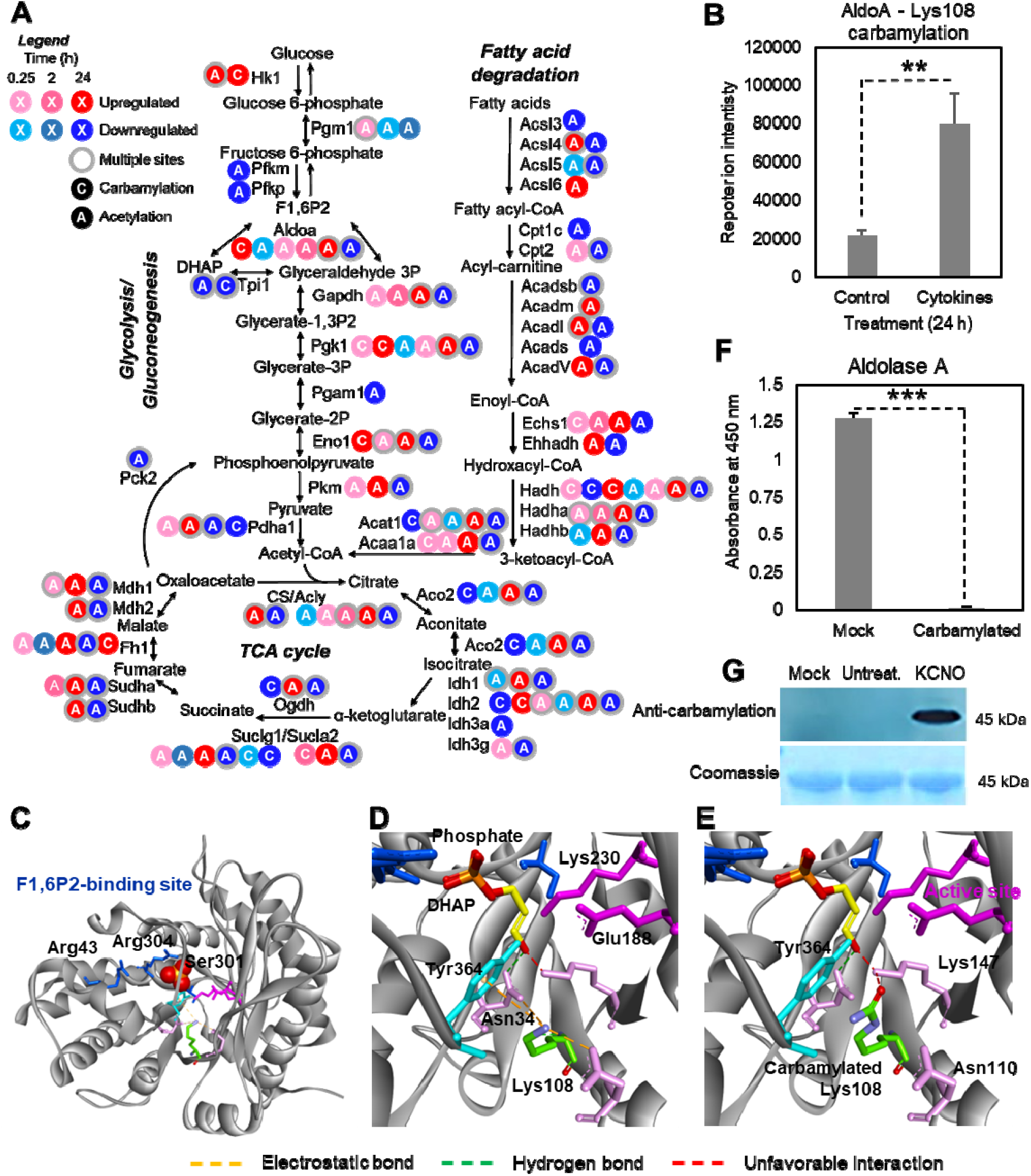
Regulation of carbon metabolism by carbamylation. (A) A map of the differentially carbamylated and acetylated proteins of the central carbon metabolism, including the Glycolysis/Gluconeogenesis, TCA cycle and Fatty acid metabolism pathways. (B) Quantification of aldolase carbamylation site at lysine 108 in the proteomics analysis. (C-E) Structure of the human aldolase A (PDB ID: 6XMH model with dihydroacetone phosphate (DHAP) from 2QUV). Lysine 108 has an electrostatic interaction with tyrosine 364 (D), which would be destroyed with carbamylation (E). (F) Enzymatic activity of carbamylated aldolase A. (G) Western blot and SDS-PAGE of potassium cyanate (KCNO) treated aldolase A to induce carbamylation. ** p≤0.01, *** p≤0.001 by Student’s *t*-test.

### Carbamylation is enriched in protein synthesis and processing pathways

Carbamylation sites regulated by the cytokine treatment were also enriched in pathways related to protein synthesis and processing, including Aminoacyl-t-RNA biosynthesis and Protein processing in endoplasmic reticulum (**Figure 2E**). In the Protein processing in endoplasmic reticulum pathway, 9 proteins were differentially carbamylated with the cytokine treatment (**Figure 6A**). These included proteins such as heat shock protein Hsp90 and protein disulfide isomerase Pdia3 with multiple regulated carbamylation sites (**Figure 6A**). An additional 26 proteins were differentially acetylated (**Figure 6A**), showing that this pathway is highly modified in response to the cytokine treatment.

**Figure 6.**
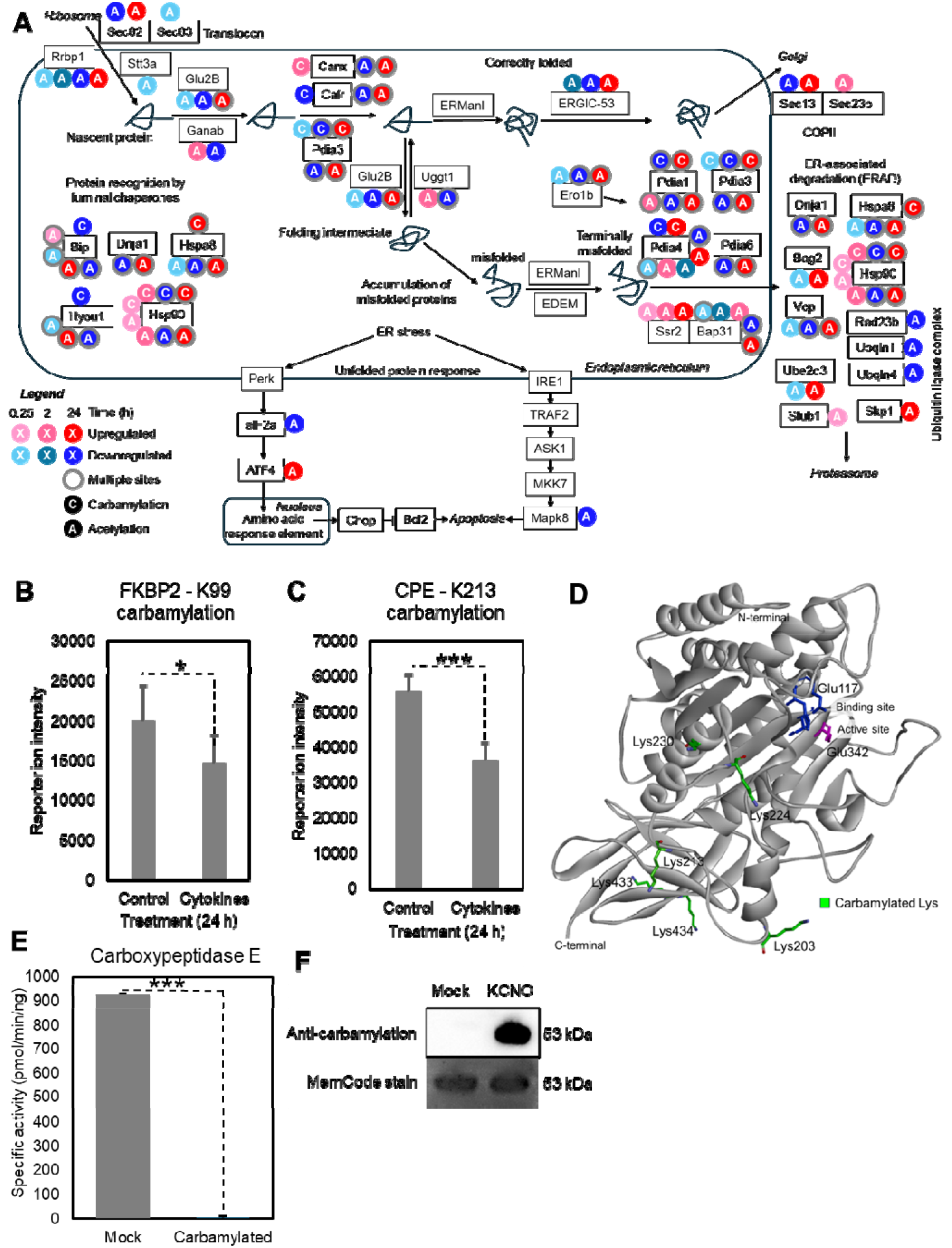
Regulation of the Protein processing in endoplasmic reticulum pathway by carbamylation. (A) A map of the differentially carbamylated and acetylated proteins of the Protein processing in endoplasmic reticulum pathway. Quantification of FK506-binding protein 2 (FKBP2) carbamylation at lysine 99 (B) and carboxypeptidase E (CPE) carbamylation at lysine 108 (C) in the proteomics analysis. (D) Structure of carboxypeptidase E highlighting carbamylation sites found in MIN6 cells (203 and 213) and human islets (K203, K224, K230, K433 and K434). (E) Enzymatic activity of carbamylated carboxypeptidase E. (F) Western blot and SDS-PAGE of potassium cyanate (KCNO) treated carboxypeptidase E to induce carbamylation. * p≤0.05, *** p≤0.001 by Student’s *t*-test.

We observed 4 insulin-processing enzymes among the proteins differentially carbamylated in response to the cytokine treatment: protein disulfide isomerases Pdia1 and Pdia4, FK506-binding protein 2 (FKBP2) and carboxypeptidase E (CPE). Pdia1 and Pdia4 were differentially carbamylated at multiple sites (**Figure 6A**), while FKBP2 had its carbamylation site at lysine 99 reduced by 27% 24-h post-cytokine treatment (**Figure 6B**). CPE was carbamylated at K203 and K213, with the latter site reduced by 35% 24-h post-cytokine treatment (**Figure 6C**). To investigate whether carbamylation of CPE occurs *in vivo*, we reanalyzed the single human islet proteomics data by Swensen et al. [40]. We identified 6 carbamylated bearing modifications at K203, K224, K230, K433, and K434, demonstrating that CPE carbamylation occurs *in vivo*. The carbamylation sites were spread in different regions of the CPE based on the crystal structure of the human ortholog (PDB ID: 2BO9) (**Figure 6D**). To determine possible functions of lysine carbamylation in regulating CPE activity, we pan-carbamylated this enzyme with potassium cyanate and measured its activity with a fluorescence assay. Carbamylation abolished CPE activity (**Figure 6E-F**). These results show that carbamylation of proteins involved in protein synthesis and processing is regulated by cytokines, and that carbamylation can affect protein processing by inhibiting enzymes like CPE.

## Discussion

Technical challenges have historically limited our understanding of the role of the PTM carbamylation. Here, we applied a novel carbamylation analysis pipeline to study possible roles of protein carbamylation in T1D. Our results identify that carbamylation is increased in islets of mice that develop T1D and in β cells treated with pro-inflammatory cytokines. Pathway enrichment identified changes in multiple pathways related to T1D development, and functional analysis confirmed direct effects on function of proteins vital for β-cell metabolism and insulin secretion. In aggregate this work suggests that this PTM could contribute to beta cell immunogenicity and dysfunction in the context of T1D.

Current knowledge of carbamylation’s roles in T1D is limited to increased carbamylation of plasma lipoproteins and hemoglobin due to poor glycemic control and nephropathy [45, 46]. Carbamylation is better characterized in rheumatoid arthritis, in which it plays a role in disease pathology. In this disease, carbamylation is induced by myeloperoxidase, which converts thiocyanate into cyanate and modifies histones. The autoimmune response induces the neutrophils to release the modified histones into neutrophil extracellular traps, turning them into neoantigens for autoantibody production [22]. These anti-carbamylation autoantibodies are pathogenic, inducing tissue damage and aggravating the arthritis symptoms [22]. Our analysis identified increased carbamylation in islets of NOD mice with insulitis. Although autoantibodies are common in T1D, anti-carbamylation autoantibodies have not yet been reported. Our pathway analysis supports the idea that this PTM may be contributing to β-cell immunogenicity, as cytokine-regulated carbamylations were enriched in proteins from pathways related to autoimmune diseases, including antigen presentation. This pathway has been proposed to be regulated mainly by transcriptional regulation [47]. However, the large extent of carbamylation and acetylation sites regulated by cytokines suggests that post-translational modifications might also play a role in regulating this pathway.

Our data also show that cytokine treatment regulates chromatin remodeling proteins including histones and acetyltransferases. In addition to the myeloperoxidase-mediated carbamylation [22], histones also have been shown to be carbamylated by carbamoyl-phosphate produced by a nuclear carbamoyl-phosphate synthase I [13]. Given the fact that carbamylation is structurally analogous to acetylation, it is reasonable to speculate that they have similar functions. However, there is currently no information in the literature on whether histone carbamylation can regulate gene expression like acetylation does. Another potential regulation point is the carbamylation of histone acetyltransferases p300 and CREBBP. For instance, p300 has been shown to be regulated by autoacetylation, including at lysine 1542 [48], the human ortholog residue of the mouse carbamylated lysine 1541. This may represent a potential crosstalk point through which carbamylation may regulate downstream histone acetylation. However, additional work is needed to test this hypothesis.

The central carbon metabolism also was enriched with proteins differentially carbamylated in response to the cytokine treatment. We showed that carbamylation of aldolase A at lysine 108 increases 4-fold with the cytokine treatment. This lysine residue is located close to the active site and interacts with tyrosine 364, an amino acid residue that helps accommodate the substrate [44]. Tyrosine 364 is essential for enzymatic activity, and nitration of this residue impairs aldolase A activity [49]. Pan-carbamylation of aldolase reduced its activity to trace levels. The inhibition of aldolase A by carbamylation is like the regulation of the central carbon metabolism enzymes by acetylation. For instance, acetylation of aldolase A at lysine 13 reduces its enzymatic activity [50]. Mechanistically, we have shown that acetylation inhibits the central carbon metabolism enzymes in bacteria by modifying and eliminating the positive charge of evolutionarily conserved lysine residues required for enzymatic activity or co-factor binding [51]. Here, we show a similar mechanism in carbamylation, in which a lysine residue that helps guide substrate binding in aldolase A is carbamylated. The fact that many enzymes are carbamylated and acetylated suggests an additive inhibitory effect on their activities.

Lastly, we found that cytokine-regulated carbamylation was enriched in proteins involved in protein synthesis and processing pathways. In Protein processing in the endoplasmic reticulum pathway, several insulin-processing proteins were differentially carbamylated. This observation could be consistent with known post-transcriptional impacts of inflammatory cytokines in the context of islet endoplasmic reticulum stress. Intriguingly, among the insulin-processing proteins, we found that CPE carbamylation at lysine 213 is downregulated by cytokine treatment and that CPE carbamylation inhibited its enzymatic activity. CPE levels are reduced in islets in T1D [52]. Therefore, the reduction in carbamylation in response to cytokine treatment may represent a compensatory mechanism to improve insulin processing and ultimately secretion. Importantly, we found multiple carbamylation sites on CPE in human islets, suggesting that this modification occurs *in vivo*. However, additional studies will be needed to determine the dynamics of carbamylation *in vivo*.

In summary, here we show that carbamylation is higher in islets of NOD mice and the levels of carbamylation regulated by cytokines in MIN6 insulin-producing cells. Carbamylation modifies proteins from a variety of pathways and may represent a regulatory mechanism for cell metabolism and insulin processing.

### Limitations of the study

A limitation of this study is that the carbamylation of enzymes was chemically induced to pan-modify them. Therefore, the contribution of individual carbamylation sites to regulating enzymatic activity still needs to be determined. With advances in genetic code expansion [53], we believe that expressing proteins with carbamylation at a specific lysine residue will be possible in the near future.

## DATA AVAILABILITY STATEMENT

Mass spectrometry raw data was deposited in MassIVE repository, a member of the ProteomeXchange Consortium, under the accession number: MSV000101719.

## ACKNOWLEDGMENTS

The authors thank Dr. Ethan Stoddard for critically reading the manuscript. Part of the work was performed in the Environmental Molecular Sciences Laboratory, a U.S. Department of Energy (DOE) national scientific user facility at Pacific Northwest National Laboratory (PNNL) in Richland, WA. Battelle operates PNNL for the DOE under contract DE-AC05-76RLO01830. This work was supported by the National Institute of Diabetes and Digestive and Kidney Diseases grants U01 DK127505 (to L.S. and E.S.N.), R01 DK138335 (to E.S.N. and B-J.M.W-R.), R01 DK133881 (to E.K.S. and R.G.M.), U01 DK127786 (to R.G.M. and E.S.N.), R01 DK121929 (to E.K.S.), and R01 DK060581 (to R.G.M.).

## AUTHOR CONTRIBUTIONS

Conceptualization: YY, ESN. Investigation, data curation, formal analysis and validation: all authors. Visualization: YY, PD, ESN. Methodology: YY, HK, MGU, MG, HEW, FL, WJQ, CVR, LS, ESN. Project administration, supervision and resources: WJQ, GC, GM, RGM, BJMWR, AJW, CVR, EKS, LS, ESN. Funding acquisition: RGM, BJMWR EKS, LS, ESN. Writing – original draft: YY, ESN. Writing – review & editing: all authors.

## Supplemental Material

### Antibody specificity

Carbamylation, acetylation and citrullination are three post-translational modifications with very similar structures (**Figure S1A**), often leading to cross-reactivity of antibodies against them [30, 54]. Therefore, we tested 4 antibodies for their specific in recognizing carbamylation. While all tested antibodies recognized carbamylation the goat polyclonal antibody (ab175576, Abcam) also recognized FBS to a certain extent (**Figure S1B**). The rabbit (STA-078, Cell Biolabs) and the goat polyclonal antibodies also recognized citrullinated FBS (**Figure S1C**). Antibodies ab175576 and STA-078 also had the strongest reactivity to acetylated FBS (**Figure S1D**). The other two antibodies that had the least cross-reactivity were quantitatively tested by having all modified antigens on the same plate. None of these antibodies recognized citrullination, but the rabbit polyclonal (22428, Cayman) recognized acetylation and carbamylation at a similar level (**Figure S1E**). The mouse monoclonal antibody (23203, Cayman) strongly recognized carbamylation with minor binding to acetylation only in high concentrations (**Figure S1F**). In summary, out of the tested antibodies, the mouse monoclonal antibody showed the best specificity and was used for the subsequent experiments.

**Figure S1.**
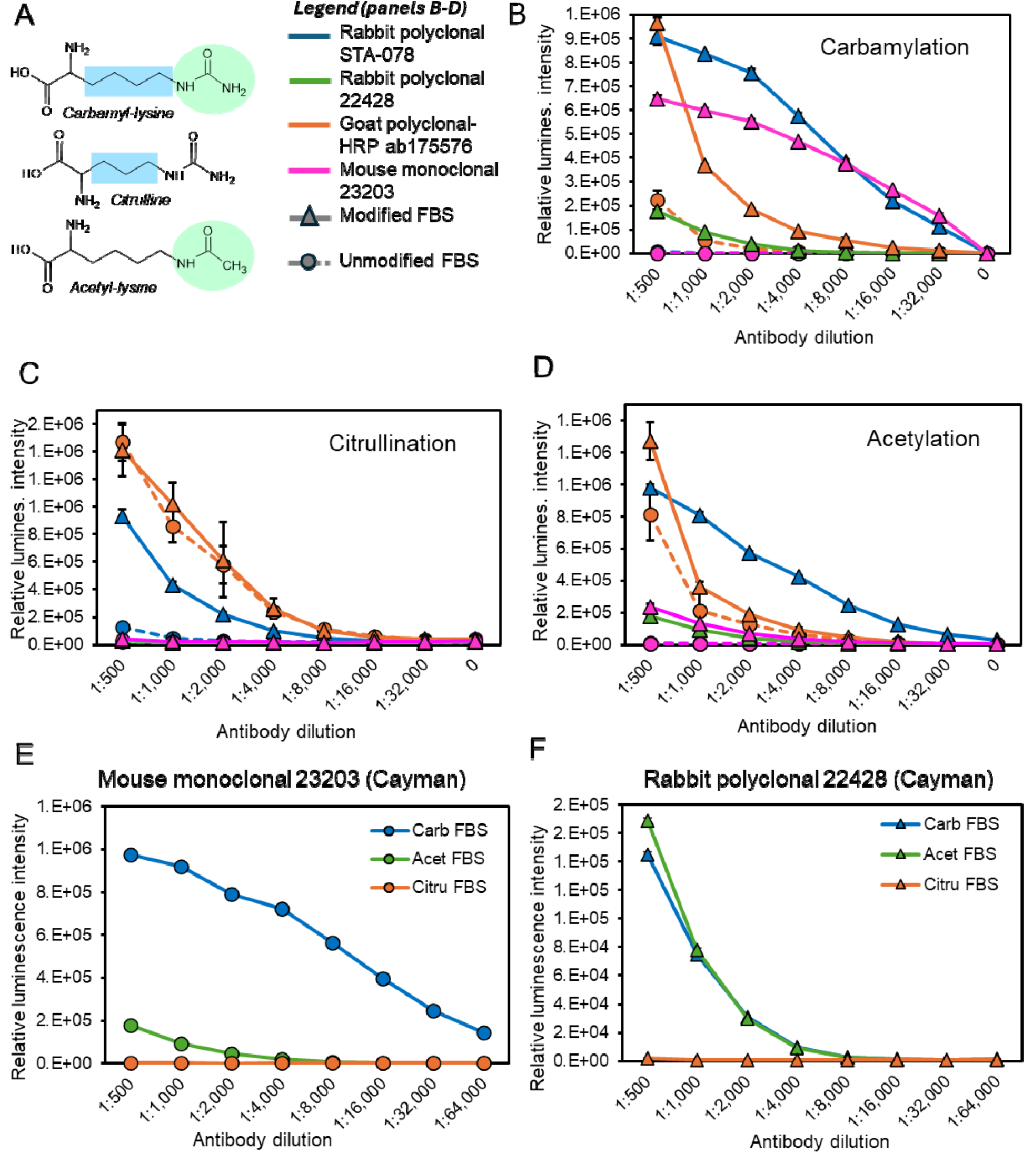
Evaluation of commercial anti-carbamylation antibody specificity. (**A**) Structure of carbamyl-lysine, citrulline, and acetyl-lysine. The differences between these amino acids are highlighted. This figure panel is adapted from You et al [11] under Creative Commons Attribution License (CC-BY 4.0). (**B-D**) Binding of antibodies to carbamylated (B), citrullinated (C) and acetylated (D) fetal bovine serum measured by chemiluminescence ELISA. (**E-F**) Binding of mouse monoclonal (E) and rabbit polyclonal (F) antibodies against carbamylated, citrullinated and acetylated fetal bovine serum.

**Figure S2.**
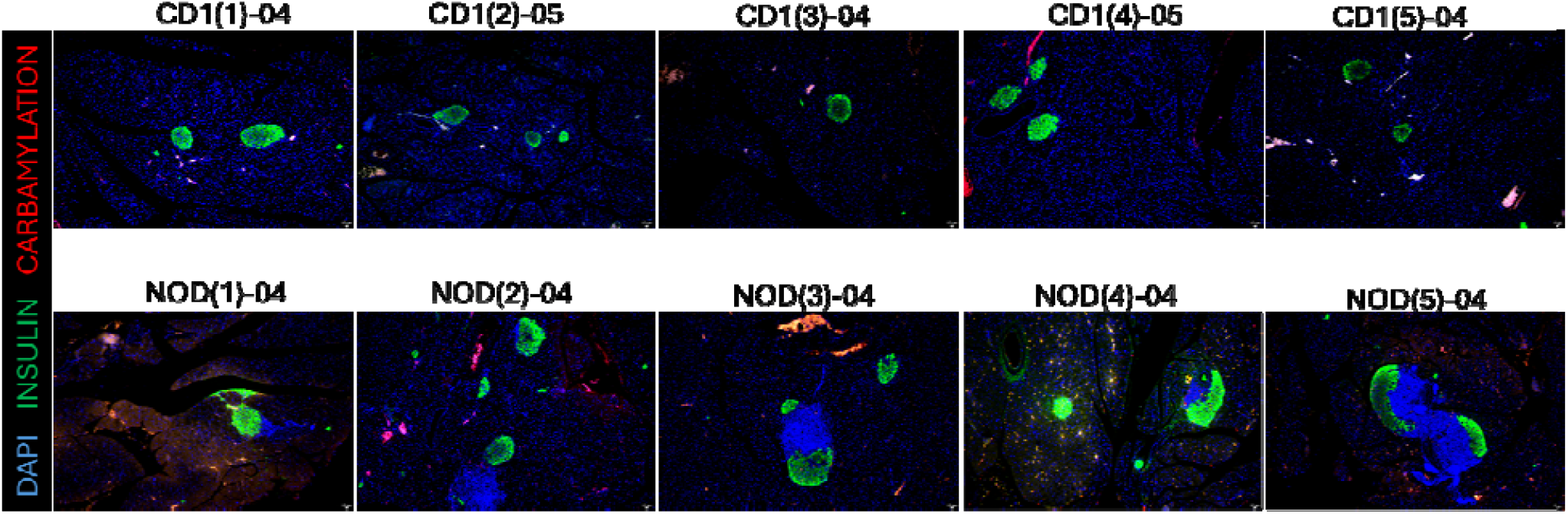
Lysine carbamylation profiles across individual non-diabetic CD1 and non-obese diabetic (NOD) mice. Pancreata from 8-9-weeks old mice were immunostained for carbamylation, insulin and nucleic acid (DAPI) and imaged in a fluorescence microscope. In NOD mice tissue, there is a noticeable infiltration of lymphocytes, marked by the high density of nuclei (DAPI staining).

